# Ecological effects of genome size in yellow starthistle (*Centaurea solstitialis*) vary between invaded and native ranges

**DOI:** 10.1101/2022.10.25.513778

**Authors:** F. Alice Cang, Katrina M. Dlugosch

## Abstract

Invasive species have become a pervasive threat on every continent and across a broad array of environments. Several traits predicted to promote invasion success, such as small seed size, rapid vegetative growth and short time to reproduction, are correlated with smaller genome sizes in a number of systems. To understand the influence of genome size on plant invasion dynamics, we compared genome sizes and traits in *Centaurea solstitialis* (YST) genotypes from the Californian invasion to those from their native source region in Spain. We conducted a common garden experiment and genome size survey to ask: (1) Is the invasion associated with genome size reduction? (2) To what extent can differences in genome size explain previously observed increases in plant size and reproduction in YST invasions? (3) Finally, we tested for expected evolutionary patterns in genome size across populations, including evidence of selection favouring reduced genome sizes at higher elevations, and evidence of stochastic processes leading to increases in genome sizes where effective population sizes are smaller. We found a reduction in corrected genome size in the invaded range, as well as significant interaction effects of range x elevation on genome size, and range x genome size on flowering time variation. Specifically, larger genomes tended to flower later and genome size decreased with increasing elevation in the invasion only. These emergent relationships in invading YST suggest potential selection for smaller genomes following introduction of YST to its invaded range. We also found a significant effect of measurement date on genome size estimation by flow cytometry, and this effect was more pronounced among native range genotypes.

## Introduction

Invasive species have become a pervasive threat on every continent and across a broad array of environments (Vitousek *et al*. 1996; Wilcove *et al*. 1998; Sala *et al*. 2000). As introduced species encounter novel environments during colonization and range expansion, they are expected to encounter novel selection pressures for traits and trait values not found in their native ranges. Indeed, there is now plenty of evidence in many systems for rapid evolution in species introductions (Prentis *et al*. 2008; Whitney and Gabler 2008). In plants, some documented rapid evolutionary changes include shifts towards selfing (Kollmann and Bañuelos 2004), emergence of clinal variation in life history traits (Dlugosch and Parker 2008; Colautti and Barrett 2013; Boheemen *et al*. 2018), altered species interactions (Sander *et al*. 2007; Seifert *et al*. 2009; Rotter and Holeski 2008), variable propagule size (Hierro *et al*. 2020), hybridization (Whitney *et al*. 2006) and structural variation in the genome such as polyploidy (Beest *et al*. 2012) and transposable element activity (Stapley *et al*. 2015). Importantly, these rapid evolutionary changes in introduced populations suggest opportunities for adaptation to play a critical role in facilitating invasions.

There has been considerable research effort in determining traits of successful invaders that might allow them to establish and spread aggressively into new environments (Liao *et al*. 2008; Dawson *et al*. 2011; Castro-Diez *et al*. 2014; Van Kleunen *et al*. 2015; Montesinos 2021). Baker (1965) was first to propose a set of characteristics of an ideal plant invader. These included traits that facilitated higher dispersal rates (such as small seed size and wind pollination), rapid vegetative growth and short time to reproduction (Baker 1965). While studies about traits of plant invaders have empirically demonstrated that invasive plants tend to have faster growth, more seeds, and more efficient resource consumption than their native counterparts (Van Kleunen *et al*. 2015), relatively few generalisations can be drawn, as selective advantages conferred by specific traits or trait values are likely to be context-dependent (Funk 2013; Kueffer *et al*. 2014). Several of these predicted “ideal weed” traits correlate with genome size (Grotkopp *et al*. 2004; Schmidt and Drake 2011; Fridley and Craddock 2015; Meyerson *et al*. 2020), which has prompted the hypothesis that reduced genome sizes may evolve in association with successful colonization events (Knight *et al*. 2005; Hidalgo *et al*. 2015; Carta and Peruzzi *et al*. 2016). Comparative analyses of genome size variation across plants have identified effects of larger genome sizes on increased cell size, longer cell cycles, decreased stomatal density and greater seed mass (Beaulieu *et al*. 2007a; Beaulieu *et al*. 2008). Larger genomes also trend towards longer minimum generation times (Grotkopp *et al*. 2004), slower photosynthetic rates (Knight *et al*. 2005) and larger plant sizes (Carta and Peruzzi 2016), suggesting cellular effects also scale to the level of entire individuals and influence ecologically relevant traits. These trait differences between individuals with large and small genomes suggest potential costs associated with maintaining large genomes, potentially constraining growth, reproduction or dispersal. All else being equal, large genomes should incur higher costs of DNA replication and require larger cell sizes.

Environmental correlates of genome size identified in several studies offer potential examples of selective effects on genome size, including temperature seasonality in *Helianthus* species (Qiu *et al*. 2018), plasticity in *Phragmites australis* (Meyerson *et al*. 2020), and annual temperature and precipitation in the giant genomes of *Lilium* species (Du *et al*. 2017). Among studies of clinal variation in genome size, special consideration has been given to effects of elevation on genome size. Smaller genomes in higher elevation populations, owing to potential differences in growing season and the fitness effects of development time, were found in maize (Bilinski *et al*. 2018), *Corchorus olitorius* (Benor *et al*. 2011), *Lagenaria siceraria* (Achigan-Dako *et al*. 2008), as well as a small number of non-crop systems such as *Dactylis glomerata* (Creber *et al*. 1994), and *Arachis duranensis* (Temsch and Greilhuber 2001). Variable environments that influence transposable element content might be mediated by interactions between environment and genome size. For example, the number of copies of the retrotransposon BARE-1 varies with elevation in wild barley accessions and its establishment on north-facing or south-facing slopes (Kalendar et al 2000). Populations from south-facing slopes have larger genomes than those from north-facing slopes, and loss of the BARE-1 copies through recombination is negatively correlated with elevation and dryness of the habitat (Vicient et al 1999; Kalendar et al 2000). However, evidence is mixed and several studies of intraspecific genome size variation fail to recover any clinal variation in genome size (Lysak *et al*. 2000; Oney-birol and Tabur 2018; Savas *et al*. 2019). This body of work reflects increasing recognition that genome size variation itself may function as a target of selection within plant populations. If genome size imposes functional constraints, a new colonist may benefit from reduced genome size effects on a suite of traits that promote the growth of its lineage, such as fast generation times and high reproductive rates. This may account for the observation that, in a survey of over 19 000 plant species across the continental United States, plants with small genome sizes were relatively over-represented among weedy and invasive taxa (Kuester et al. 2014). However, understanding of the population consequences and fitness effects of these associations is currently limited (Gaut and Ross-Ibarra 2008; Whitney *et al*. 2010; Mei *et al*. 2017; Bilinski *et al*. 2018), and few studies have examined how these effects influence invasion dynamics (although see review in Suda *et al*. 2014).

The opportunities for genome size to contribute to colonization success are particularly strong in plants, which demonstrate the greatest breadth of genome size variation among major groups of eurkaryotes (Gregory 2005). This variation spans four orders of magnitude within flowering plants alone, from *Genlisea aurea* with the smallest known genome size (63 MB) (Leushkin *et al*. 2013) to *Paris japonica* (148 GB) with the largest eukaryotic genome recorded to date (Pellicer *et al*. 2010). Although historically controversial (Greilhuber 2005), the lability of plant genome size is also found at smaller phylogenetic scales among individuals within species (Zaitlin and Pierce 2010; Bilinski *et al*. 2018; Stelzer *et al*. 2019; Boutte *et al*. 2020). Increasingly, the effects of colonization on intraspecific genome size variation have been examined by comparing populations in the native and introduced ranges (Crosby *et al*. 2014; Meyerson *et al*. 2020). For example, populations of *Avena barbata*, an invasive oatgrass, demonstrated smaller median genome sizes from the invasive Californian accessions than those from the native range (Crosby et al. 2014). However, studies that can draw direct comparisons between an invasive population and its source region are scarce, and issues of methodology in estimating genome size can produce variable results (Lavergne *et al*. 2009, Martinez *et al*. 2018).

Critically, colonization of new environments necessarily exposes the introduced species to novel stressors that can break the mechanisms maintaining genome size. Transposable element dynamics strongly influence genome size in plants (Tenaillon *et al*. 2010), and host genome control of transposable element proliferation is sensitive to environmental stress (Grandbastien 1998; Chenais *et al*. 2012; Horvath *et al*. 2017). An association between transposable element activity and stressful environments is often considered evidence of their mobilisation as an agent of adaptation (Chenais *et al*. 2012; Casacuberta and Gonzalez 2013; Negi *et al*. 2016). However, as with any other mutation, most transposable element mobility is likely to be selectively deleterious or neutral (Tenaillon *et al*. 2010; Pasyukova *et al*. 2004; Lockton *et al*. 2008). Analysis of the *Arabidopsis lyrata* and closely related *A. thaliana* genomes reveals that the larger genome of *A. lyrata* bears signatures of ongoing selection for DNA loss (Hu *et al*. 2011). Long terminal repeats, a retrotransposon thought to be a primary driver of genome size differences in plants, are typically younger than 3 million years old across taxa (Baidouri and Panaud 2013), which suggests that DNA purging is common and most elements are quickly lost to purifying selection following insertion (Michael 2014). If effective population sizes are small or selection is insufficiently strong, populations may accumulate larger genome sizes due to low efficacy of purifying selection against these repetitive genetic elements (Baduel *et al*.2019).

Here, we investigate genome size variation across the native and invaded ranges of the invasive species yellow starthistle, *Centaurea solstitialis* L., in the Asteraceae (hereafter, YST). YST is an annual, obligately outcrossing, diploid thistle (2*n*=16, Heiser and Whitaker 1948; Widmer *et al*. 2007; Öztürk *et al*. 2009), and previous work has suggested that genome size is variable within its invaded ranges (Irimia *et al*. 2017). YST likely originated in the eastern Mediterranean preceding an ancient expansion into Eurasia and western Europe (Barker *et al*. 2017). Historical records indicate that it was introduced to South America from Spain and to the US from Chile, via imports of alfalfa, with the earliest recorded North American occurrence in the San Francisco Bay area in 1869 (Pitcairn *et al*. 2006). This recent colonization history has been confirmed by genetic studies (Dlugosch *et al*. 2013; Eriksen *et al*. 2014; Barker *et al*. 2017). Since its introduction into North America in the 19th century, YST has expanded its range in all directions across the western United States, and is particularly problematic in California (Maddox and Mayfield 1985; Pitcairn *et al*. 2006). We know from previous work that Californian YST populations are derived from source populations found in western Europe (Dlugosch *et al*. 2013; Barker *et al*. 2017), have evolved shifts in growth rate and flowering time (Dlugosch *et al*. 2015), and have expanded across large elevational and environmental gradients (Gerlach 1997; Pitcairn *et al*. 2006; Braasch *et al*. 2019). Importantly, severe YST invasions in California are associated with the evolution of increased size and earlier development, leading to large increases in reproduction capacity (Dlugosch *et al*. 2015). These recent changes in genome size-associated traits provide an opportunity to examine relationships between genome size and rapid trait evolution of a plant invader following major colonization events.

In this study, we compared genome sizes and traits of YST genotypes from the Californian invasion to those from their native source region in Spain. We asked whether invasion was associated with genome size reduction, and we investigated to what extent differences in genome size explained previously observed increases in plant size and reproduction in YST invasions. Finally, we tested for expected evolutionary patterns in genome size across populations, including evidence of selection favouring reduced genome sizes at higher elevations, and evidence of stochastic processes leading to increases in genome sizes where effective population sizes are smaller.

## Materials and Methods

### Common garden

Seeds were collected from four sites in the invaded range of California, USA in August 2016, and from four sites in the native range in Spain in September 2008 (Table 1). The four sites in the invaded range were selected to reflect the oldest areas of introduction in North America, near the San Francisco Bay area, which has the earliest records of introduction (Gerlach 1997; Braasch et al. 2019). At each site, fruiting capitula were collected along a linear transect, from maternal plants >1m apart. We grew 20-25 individuals from each of the four invaded range populations, and 3-17 individuals from each of the four native range populations (N=119). We germinated seeds on the surface of moist potting soil [3:2:1 ratio of Sunshine Mix #3 soil (Sun Gro Horticulture, Canada), vermiculite, and 20 grit silica sand] under fluorescent lights and 12 hour days in December 2016 and recorded germination date daily. We germinated multiple seeds from the same maternal plant and transplanted the first germinating individual at 5 weeks into 410ml Deepots (Steuwe and Sons, Inc, USA) in January 2017. Seedlings were grown in a University of Arizona greenhouse in Tucson, AZ, USA. After transplanting, we randomly assigned one individual from each population to each of up to 25 different blocks. Plants were watered daily using an automatic drip watering system and maintained through senescence.

**Table 1:** Field collections from the YST native range in Spain and invaded range in California, USA. Sample sizes for genome size estimates from each population include the number of individuals with at least 1 measurement. Values in parentheses indicate how many individuals out of the total have replicate estimates.

|  |  |  | Coordinates | Elevation<br>(m) | N (Common<br>garden) | N (Genome<br>size) |
| --- | --- | --- | --- | --- | --- | --- |
| Invaded<br>range | Diablo | DIA | N 37.863694 W 121.976444 | 237 | 20 (3) | 20 (3) |
|  | Gilroy | GIL | N 37.03373 W 121.53674 | 68 | 22 (4) | 22 (4) |
|  | Marin | MAR | N 38.01154 W 122.612222 | 143 | 25 (1) | 24 (1) |
|  | Red Bluff | RB | N 40.27083 W 122.271 | 157 | 23 (1) | 23 (1) |
| Native<br>range | Canales | CAN | N 41.00030 W 4.89718 | 879 | 3 (0) | 3 (0) |
|  | Cuenca | CUE | N 40.12946 W 2.13950 | 922 | 5 (0) | 5 (0) |
|  | Granada | GRA | N 37.26836 W 3.66497 | 673 | 4 (1) | 0 (0) |
|  | Salamanca | SAL | N 40.99003 W 5.65856 | 818 | 17 (1) | 16 (1) |
|  |  |  |  | Total | 119 | 113 |

Prior to bolting, we measured leaf number, length of longest leaf and width of longest leaf to estimate a size index and early growth rate (Dlugosch *et al*. 2015). We calculated a size index from leaf measurements by (leaf number*(maximum leaf length*maximum leaf width)½), which correlates linearly with biomass in this species (Dlugosch *et al*. 2015). Linear growth rates were calculated using this size index and number of days since germination. To determine time to reproduction, the common garden was checked daily to record days to the initiation of bolting and days until the first flower. Plants were harvested when most individuals had died, and we counted the final number of flowering heads. Aboveground biomass was quantified after drying overnight at 60°C.

### Genome size estimation

We estimated genome size by flow cytometry using the FACSCanto II instrument (BD Biosciences, San Jose, CA, USA) equipped with a blue (488-nm), air-cooled, 20-mW solid state laser and a red (633-nm) 17-mW helium neon laser for UV excitation. Sample preparation followed a modified two-step protocol by Doležel *et al*. (2007). We chose *Raphanus sativus* (2C=1.11pg) provided by the Institute of Experimental Botany (Prague, Czech Republic) as our internal standard, as it has a close but non-overlapping genome size with the previously reported nuclear genome size of YST (2C=1.74pg; Bancheva and Greilhuber 2006). Nuclear suspensions were prepared by chopping 50-60 mg each of YST and *R. sativus* fresh tissue with 0.5 ml of ice-cold Otto I buffer in a glass Petri dish on top of ice, using a new razor blade for each estimate. The suspension was filtered first through gauze, and then again through 18 μm Nylon mesh. Nuclear suspensions in Otto I buffer are considered relatively stable (Doležel *et al*. 2007) and samples were left covered and on ice until immediately before analysis by flow cytometry, at which point we added 0.77ml of Otto II buffer, 29 μL of 1 mg/mL propidium iodide (PI) stain and 7.5 μL of 1 mg/mL RNase. The sample was then gently vortexed and left for at least 5 minutes on ice in a covered container. YST genome sizes were calculated according to Doležel and Bartoš (2005). We recorded 3 estimates of genome size on different days for a subset of individuals (N=25), including 1-3 individuals from all 14 populations. We obtained a single estimate of genome size for the remaining individuals.

### Genome size correction

Our previous analyses of genome size in a large concurrent study of YST indicated that the date of flow cytometry measurement had a weakly negative, but highly significant, effect on estimated genome size (Chapter 1), in which flow cytometry estimates of genome size decreased as measurements were taken later in the season. This trend might reflect variation induced by developmental changes in the plant as it ages or changes in the flow cytometer over time (Doležel and Bartoš 2005; Doležel *et al*. 2007). To avoid systematic underestimates of genome size over time, we calculated a corrected genome size using coefficients determined from linear models explaining genome size variation (Table 2). A significantly negative effect of an interaction between estimation date and range indicated the degree of underestimation over time was more severe for native range genome sizes (Table 2). Specifically, the corrected genome size estimate = observed genome size estimate - (*β*_range_*days between germination and flow cytometry measurement), in which *β* is the range-specific coefficient (*β*_invaded_=−0.0003171, *β*_native_=−0.0010693). We used this corrected genome size in all subsequent trait analyses.

**Table 2.**
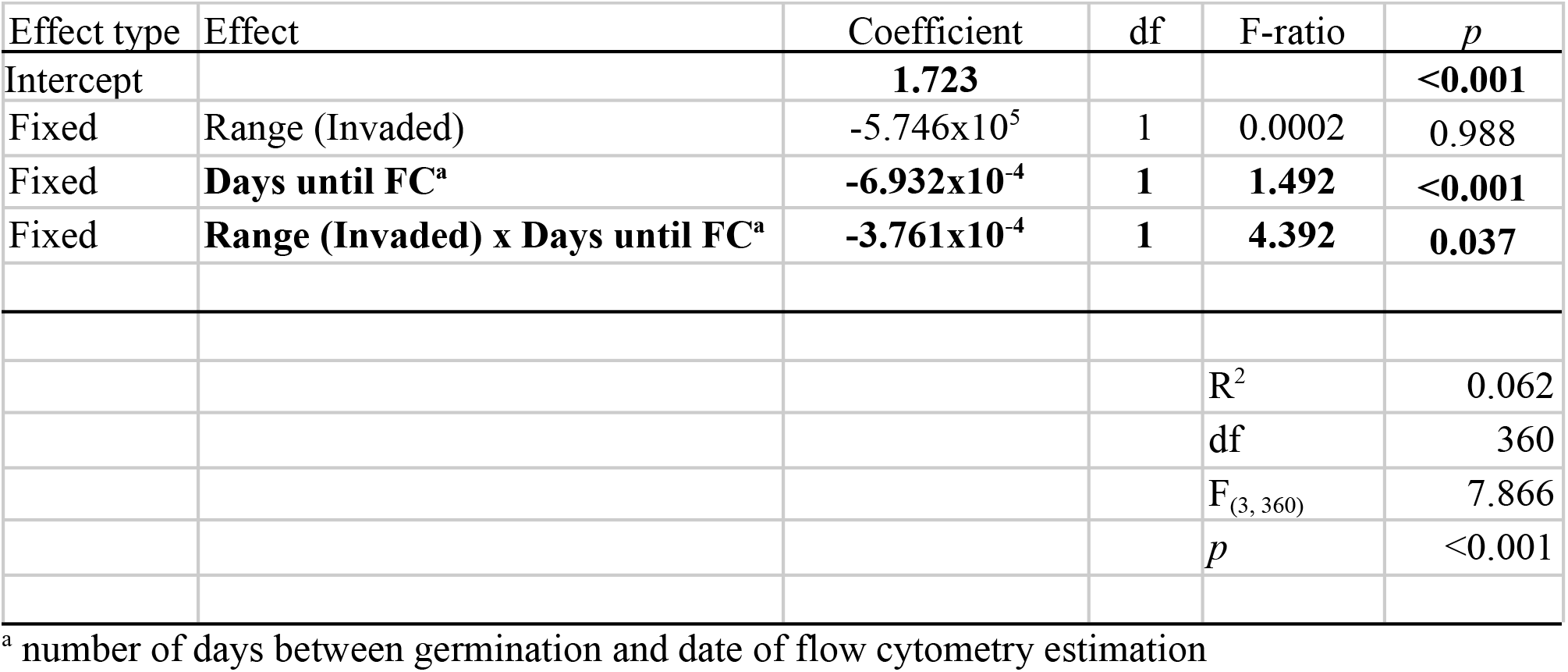
Linear model identifying the effects of date of estimation by flow cytometry on genome size estimates, for the invaded and native ranges.

| Effect type | Effect | Coefficient | df | F-ratio | *p* |
| --- | --- | --- | --- | --- | --- |
| Intercept |  | **1.723** |  |  | **<0.001** |
| Fixed | Range (Invaded) | -5.746x10<sup>5</sup> | 1 | 0.0002 | 0.988 |
| Fixed | **Days until FC<sup>a</sup>** | **-6.932x10<sup>-4</sup>** | **1** | **1.492** | **<0.001** |
| Fixed | **Range (Invaded) x Days until FC<sup>a</sup>** | **-3.761x10<sup>-4</sup>** | **1** | **4.392** | **0.037** |
|  |  |  |  | R<sup>2</sup> | 0.062 |
|  |  |  |  | df | 360 |
|  |  |  |  | F<sub>(3, 360)</sub> | 7.866 |
|  |  |  |  | *p* | <0.001 |

<sup>a</sup> number of days between germination and date of flow cytometry estimation

### Statistical analyses

We used linear models to test for (i) differences in genome size between ranges, (ii) the effects of demographic and environmental factors on genome size within and between ranges, and (iii) the effects of genome size on plant traits. Range (native vs. invaded) was included as a predictive effect within full models. Where we found significant interactions between range and other predictive variables, additional models were fit separately to each range to better explore trait and genome size variation within regions. We fit all models using *lm* in R package *stats* (R Core Team, 2021), and calculated F-ratios using *Anova* from the *car* package (Fox and Weisberg 2019).

We predicted genome size using fixed effects of range, elevation, and effective population size (N_e_). N_e_ values were obtained from published estimates for the same populations by Braasch and colleagues (2019). These values were estimated using linkage disequilibrium among genome-wide markers (LD-N_e_), a metric sensitive to recent changes in effective population size (Do *et al*. 2014). The growth and flowering traits that we collected are likely to be correlated as part of plant development, and so we used a principal component analysis (PCA) to identify major axes of variation in these traits (R Core Team, 2021). Models predicting trait PC scores included several potential effects in addition to a fixed effect of corrected genome size. To account for additional trait divergence among populations and environments, we included fixed effects of range (native vs. invaded), elevation, and populations nested within ranges. To account for variation introduced by our experimental design, we included fixed effects of days until harvest (plant age) and greenhouse position. In our previous analyses (see Chapter 1), we observed a significant effect of greenhouse location on growth, wherein individuals with smaller genomes grew more slowly and produced less biomass overall at positions closer to our greenhouse cooling system. Here, we treat greenhouse location, assigned a score from 1-9 where position 1 is coolest, as a continuous numerical variable to avoid overfitting, particularly among our native range samples.

In all cases, we fitted a full model with all effects and manually dropped non-significant effects and interactions. Higher order interactions (ie. more than two-way interactions) were excluded from full models to avoid overfitting. To select the best-fitting model among competing linear models, we competed models using *anova* in base R where models were nested, and R package *lmtest* for non-nested models (Zeileis and Hothorn 2002). In all cases, we preferred a reduced model if it was not a significantly worse fit from a model with more predictive terms.

## Results

Our survey of time-corrected genome sizes across populations in Spain and California identified intraspecific genome size variation of 1.24-fold across all individuals, ranging from 1.610-1.992pg (1.570 - 1.948 GB) with an overall mean of 1.811pg (1.771GB) (Fig. 1). The distribution overall appears bimodal, with peaks at the invaded range mean of 1.775pg (1.736 GB) and native range mean of 1.922pg (1.880 GB) (Fig. 1). Time-corrected native range genome sizes are significantly larger than invaded range genome sizes, ranging between 1.840-1.992pg (1.80-1.95 GB) and 1.61-1.88pg (1.57-1.84 GB), respectively (t = −18.662, df = 51.785, p-value<0.001) (Fig. 1, Fig. 2a). Uncorrected observed genome sizes do not differ significantly between ranges (t=−0.936, df=43.946, p=0.355, Fig. S1). The best-fitting linear model explaining variation in corrected genome sizes across both ranges identified a significant negative association with elevation only in the invaded range (Table 3, Fig. 4b-d). Elevation did not explain variation across ranges, or in the native range alone (Table 3, Fig 4b,d). We found no significant relationships between genome size and effective population size at any geographic scale (Table 3).

**Table 3.**
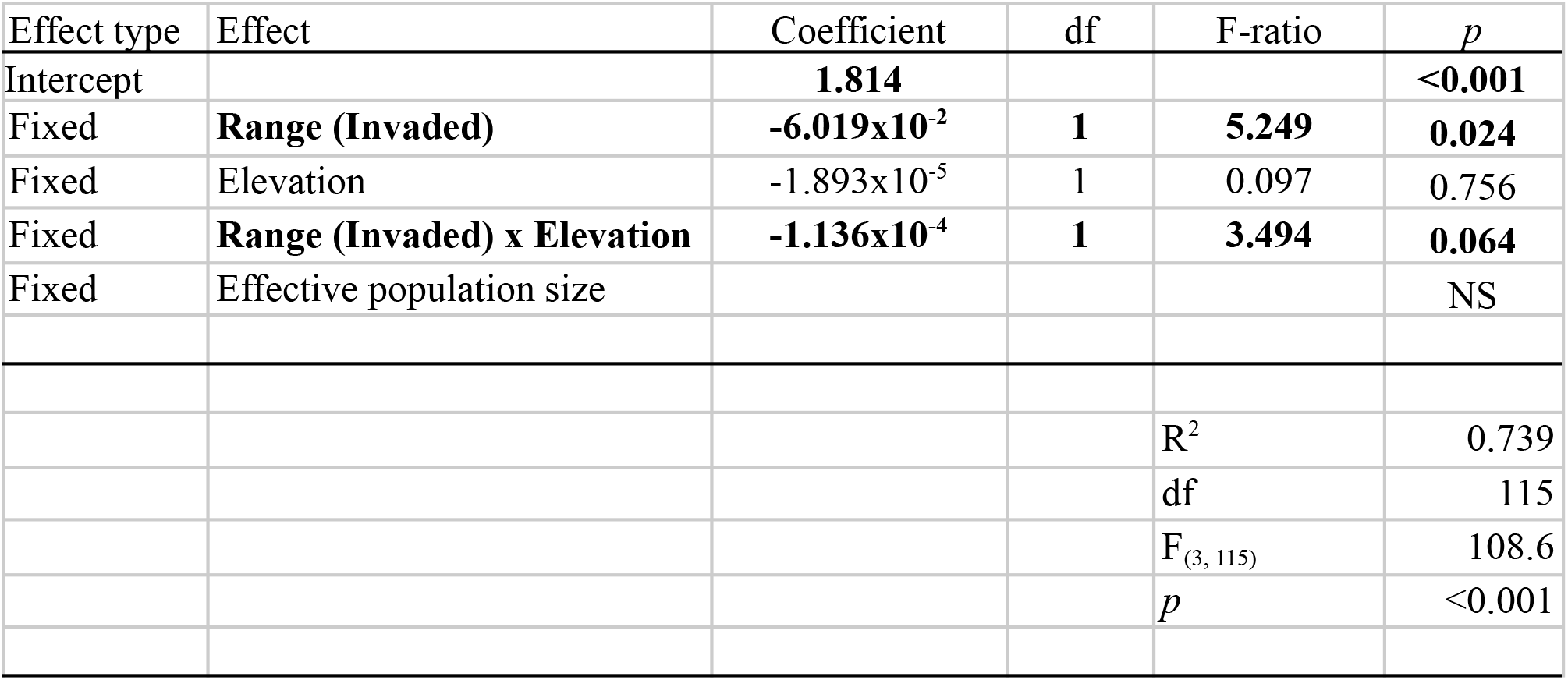
Linear model explaining variation in genome size across native and invaded range populations after correcting for estimation date. Significant effects are shown in bold. Effects without significant main or interaction effects (*p* >0.1) were removed from the model (NS).

**Figure 1:**
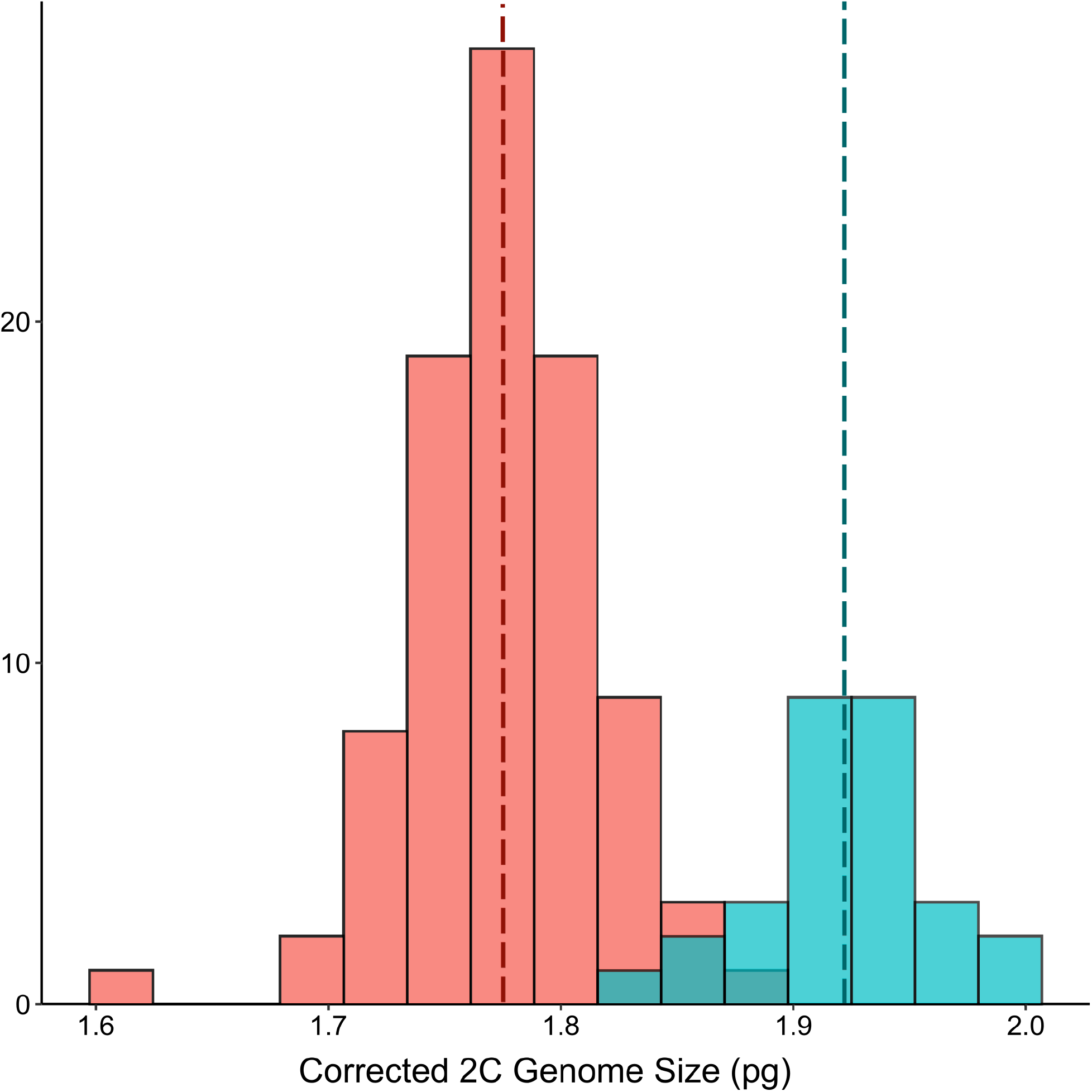
Histogram of corrected genome sizes from native range populations (blue) and invaded range populations from coastal California (pink), after range-specific corrections for estimation date. The dashed vertical lines indicate the mean genome size in each range(native range:1.922pg (N=29); invaded range: 1.775pg (N=90); mean across both ranges: 1.811pg (N=119).

**Figure 2:**
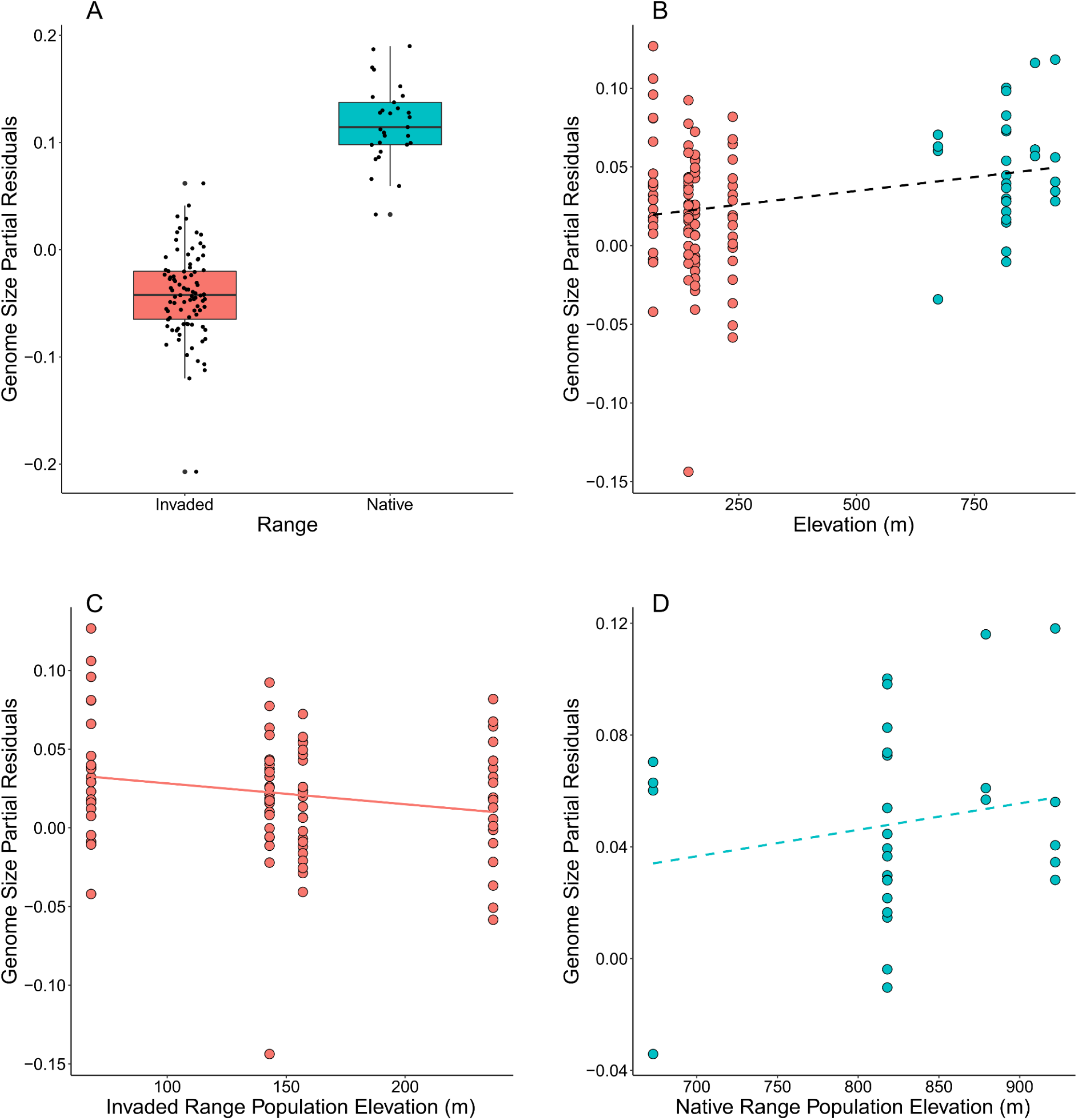
Partial residual plots from the best-fitting model explaining variation in corrected genome size across native and invaded range genotypes due to (a) range, and (b-d) elevation. Partial residuals of elevation were plotted across the entire dataset (b), and separately for each range (c, d). Solid lines indicate a significant or marginally significant relationship (*p*<0.1), and dashed lines indicate the relationship is not significant.

Principal component analysis of growth and flowering traits resulted in the first two principal components explaining 73.1% of total trait variance, with PC1 and PC2 accounting for 46.8% and 25.3%, respectively (Fig. 3). PC1 was strongly associated with early growth rates and aboveground dry biomass, while PC2 was associated with time to bolting and flowering, as well as flower number at harvest (Fig 3). Higher PC1 values correlated with faster growth and larger plant size, while higher PC2 values correlated with later bolting, later flowering, and fewer capitula produced overall. In all subsequent analysis of trait variation, we used coordinate values of PC1 and PC2 as metrics of growth and development, respectively.

**Figure 3:**
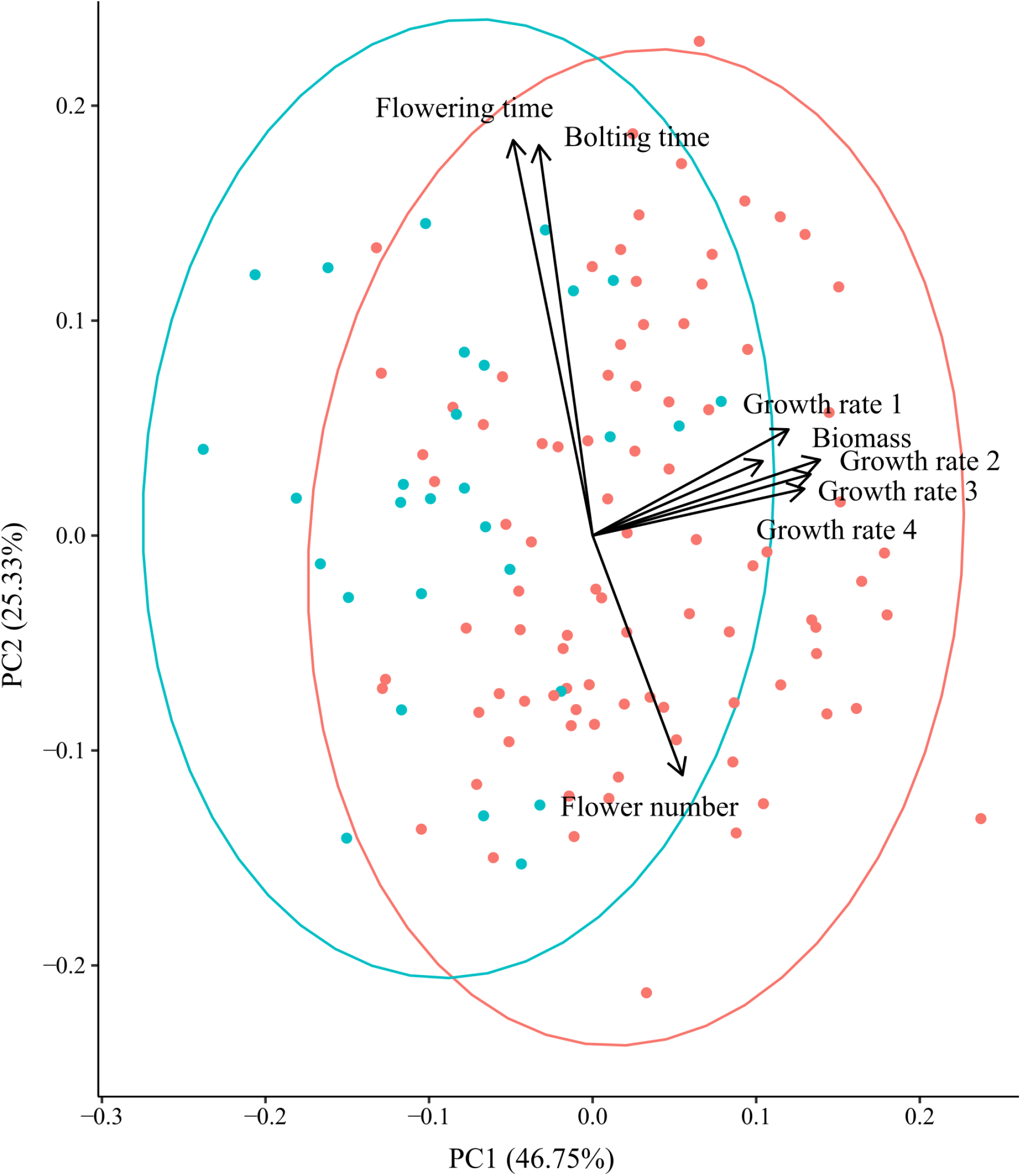
Principal component analysis of traits where each point indicates an individual plant within the common garden from native range populations (blue) and invaded range populations in California (pink) (N=119) with PC1 and PC2 explaining 72.08% of all variation. PC1 is strongly correlated with growth traits and PC2 is strongly correlated with development and reproduction traits.

Invasive populations had significantly higher mean PC1 values than native range populations (mean PC1, invaded range = 0.492, mean PC1, native range = −1.356), corresponding to faster early growth and greater dry biomass overall (t=4.6556, df=43.497, p<0.001). In our best-fitting linear model of PC1 variation, we identified a significant negative association between PC1 and elevation, such that lower elevation populations had faster growth rates and reached larger sizes (Table 4, Fig. 4b), and the relationship persisted within each range separately (Fig. 4c, d). Despite higher PC1 scores among California genotypes, we found that native range origins positively contributed to PC1 values (Table 4, Fig. 4a), contributing to faster early growth rates and aboveground biomass in the native range. Notably, all native range populations were sampled from higher elevations than any of the invasive populations in coastal California, and this positive effect of native range origins only appears in models explicitly accounting for these confounding effects (Table 4, Fig. 4a).

**Figure 4:**
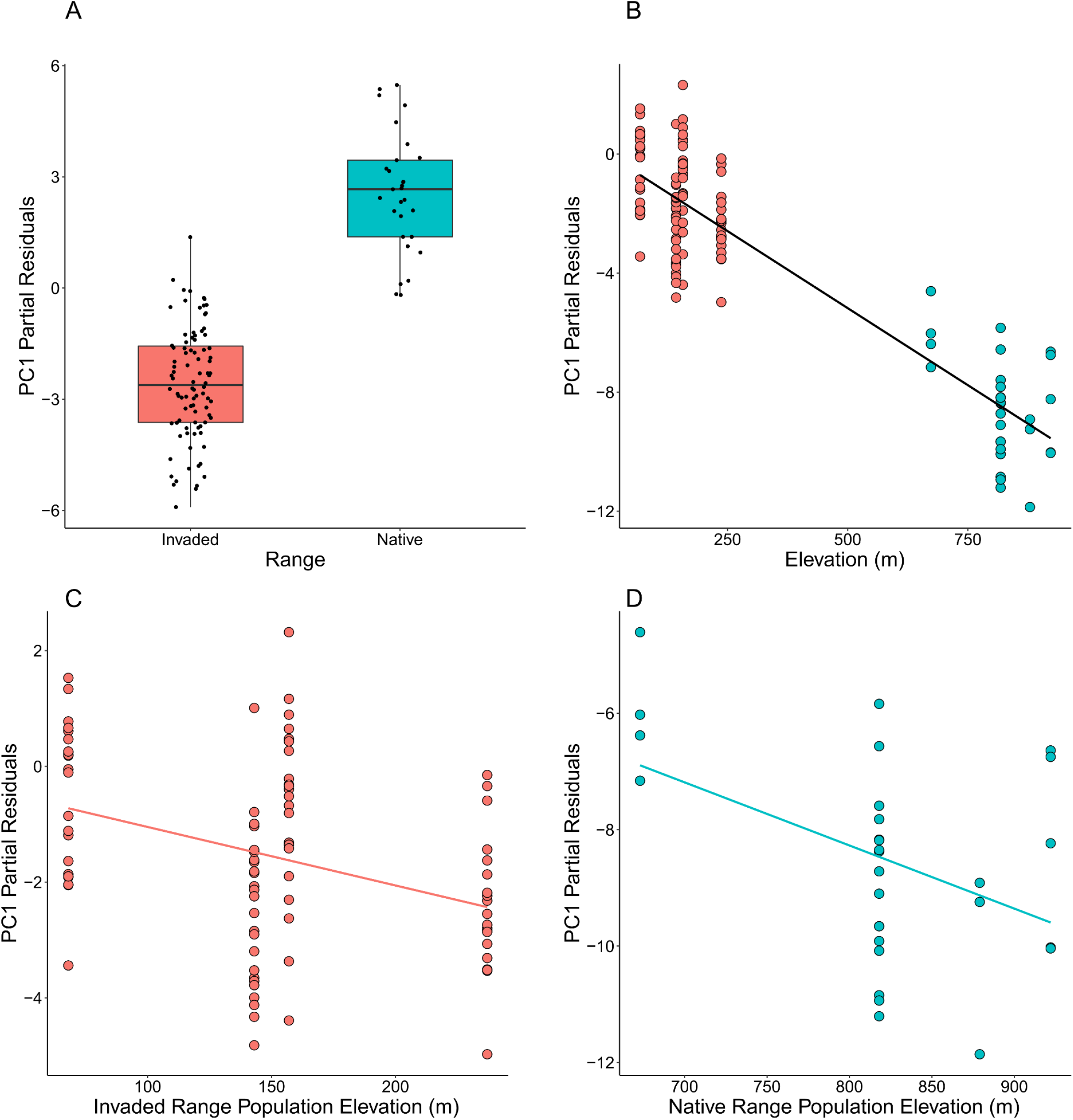
Partial residual plots from the best-fitting model explaining variation in PC1 scores across native and invaded range genotypes due to (a) range and (b-d) elevation. Partial residuals of elevation were plotted across the entire dataset (b), and separately for each range (c,d). Solid lines indicate a significant or marginally significant relationship (*p*<0.1), and dashed lines indicate the relationship is not significant.

**Table 4.** Linear model explaining PC1 (growth-related traits) variation across native and invaded range populations. Significant effects are shown in bold. Effects without significant main or interaction effects (*p* >0.1) were removed from the model (NS).

| Effect type | Effect | Coefficient | df | F-ratio | $p$ |
| --- | --- | --- | --- | --- | --- |
| Intercept |  | 1.425 |  |  | 0.713 |
| Fixed | **Range (Invaded)** | **-2.571** | **1** | **10.346** | **0.0017** |
| Fixed | **Elevation** | **-0.010** | **1** | **19.796** | **<0.001** |
| Fixed | Days until harvest | 0.013 | 1 | 0.968 | 0.327 |
| Fixed | **Greenhouse position** | **-0.169** | **1** | **9.046** | **0.003** |
| Fixed | **Days until harvest x Greenhouse pos.<sup>a</sup>** | **0.015** | **1** | **8.543** | **0.004** |
| Fixed | Corrected genome size<sup>b</sup> |  |  |  | NS |
| Fixed | Higher order interactions |  |  |  | NS |
|  |  |  |  | R<sup>2</sup> | 0.3726 |
|  |  |  |  | df | 113 |
|  |  |  |  | F<sub>(5,113)</sub> | 13.42 |
| | | | | $p$ | <0.001 |

<sup>a</sup> Includes a range-specific 2C genome size correction for effect of estimation date

In our best-fitting linear model of PC2 variation, native range origins contributed negatively to PC2, indicating faster development for native range genotypes after accounting for other fixed effects (Table 5, Fig 5a). Here, we found potential evidence for variation in selective effects of genome size on development. Although corrected genome size did not demonstrate a significant association with PC2 overall (Table 5, Fig. 5b), it interacted significantly with range (Table 5), consistent with the hypothesis that mechanisms by which large genomes impose functional constraints differ across ranges. Specifically, genome size predicted PC2 variation only in the invaded range, where it had a marginally significant positive association (Table 6, Fig 6a), but no significant relationship in the native range (Fig. 6b). Similarly, we found the nature of the relationship between elevation and PC2 also varied by range (Table 5). Among invasive genotypes, elevation had a significantly negative relationship with PC2, consistent with a scenario of accelerated development at higher elevations where the growing season is shorter (Table 6, Fig. 6c). However, this trend was reversed and significantly positive in the native range, where higher elevations were associated with longer times to reproduction (Table 7, Fig. 6d).

**Table 5.** Best-fitting linear model explaining PC2 (composite of development time and flowering traits). Significant effects are shown in bold. Effects without significant main or interaction effects (*p* >0.1) were removed from the model (NS)

| Effect type | Effect | Coefficient | df | F-ratio | $p$ |
| --- | --- | --- | --- | --- | --- |
| Intercept |  | **-2.268** |  |  | **0.009** |
| Fixed | **Range (Invaded)** | **1.612** | **1** | **3.588** | **0.061** |
| Fixed | Corrected genome size<sup>a</sup> | 5.497 | 1 | 2.318 | 0.131 |
| Fixed | Elevation | 0.001 | 1 | 0.334 | 0.565 |
| Fixed | Days until harvest | 0.011 | 1 | 1.145 | 0.287 |
| Fixed | **Greenhouse position** | **0.173** | **1** | **15.934** | **<0.001** |
| Fixed | **Range (Invaded) x Genome size<sup>a</sup>** | **6.837** | **1** | **3.598** | **0.060** |
| Fixed | **Range (Invaded) x Elevation** | **-0.007** | **1** | **14.159** | **<0.001** |
| Fixed | **Days until harvest x Greenhouse pos.** | **0.011** | **1** | **7.633** | **0.007** |
| Fixed | Higher order interactions |  |  |  | NS |
| Random | Block |  |  |  | NS |
|  |  |  |  | R<sup>2</sup> | 0.3603 |
|  |  |  |  | df | 110 |
|  |  |  |  | F<sub>(8,110)</sub> | 7.744 |
| | | | | $p$ | <0.001 |

<sup>a</sup> Includes a range-specific 2C genome size correction for effect of estimation date

**Table 6.** Best-fitting linear model explaining PC2 (composite of development time and flowering traits) across coastal invaded range populations only. Significant effects are shown in bold. Effects without significant main or interaction effects (*p* >0.1) were removed from the model (NS)

| Effect type | Effect | Coefficient | df | F-ratio | $p$ |
| --- | --- | --- | --- | --- | --- |
| Intercept |  | -0.073 |  |  | 0.563 |
| Fixed | **Genome size<sup>a</sup>** | **12.312** | **1** | **14.102** | **<0.001** |
| Fixed | **Elevation** | **-0.006** | **1** | **7.655** | **0.007** |
| Fixed | Days until harvest | 0.009 | 1 | 0.604 | 0.439 |
| Fixed | **Greenhouse position** | **0.195** | **1** | **15.549** | **<0.001** |
| Fixed | **Days until harvest x Greenhouse pos.** | **0.010** | **1** | **4.512** | **0.037** |
| Fixed | Higher order interactions |  |  |  | NS |
| Random | Block |  |  |  | NS |
|  |  |  |  | R<sup>2</sup> | 0.3783 |
|  |  |  |  | df | 84 |
|  |  |  |  | F<sub>(5,84)</sub> | 10.22 |
| | | | | $p$ | <0.001 |

<sup>a</sup> Includes a range-specific 2C genome size correction for effect of estimation date

**Table 7.** Best-fitting linear model explaining PC2 (composite of development time and flowering traits) across native range populations only. Significant effects are shown in bold. Effects without significant main or interaction effects (*p* >0.1) were removed from the model (NS)

| Effect type | Effect | Coefficient | df | F-ratio | $p$ |
| --- | --- | --- | --- | --- | --- |
| Intercept |  | **-6.47** |  |  | **0.025** |
| Fixed | **Elevation** | **0.008** | **1** | **5.850** | **0.023** |
| Fixed | Genome size<sup>a</sup> |  |  |  | NS |
| Fixed | Days until harvest |  |  |  | NS |
| Fixed | Greenhouse position |  |  |  | NS |
| Fixed | Higher order interactions |  |  |  | NS |
| Random | Block |  |  |  | NS |
|  |  |  |  | R<sup>2</sup> | 0.1781 |
|  |  |  |  | df | 27 |
|  |  |  |  | F<sub>(1,27)</sub> | 5.85 |
| | | | | $p$ | 0.0231 |

<sup>a</sup> Includes a range-specific 2C genome size correction for effect of estimation date

**Figure 5:**
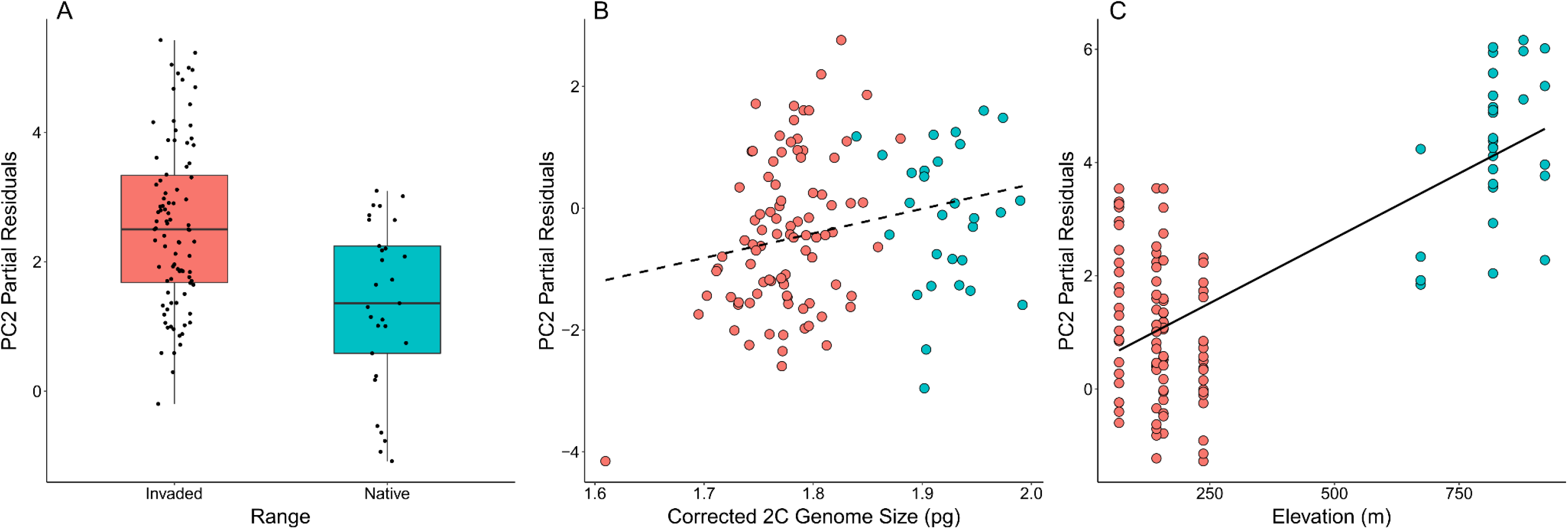
Partial residual plots from the best-fitting model explaining variation in PC2 scores across native and invaded range genotypes due to (a) range, (b) corrected genome size and (c) elevation. Solid lines indicate a significant or marginally significant relationship (*p*<0.1), and dashed lines indicate the relationship is not significant.

**Figure 6:**
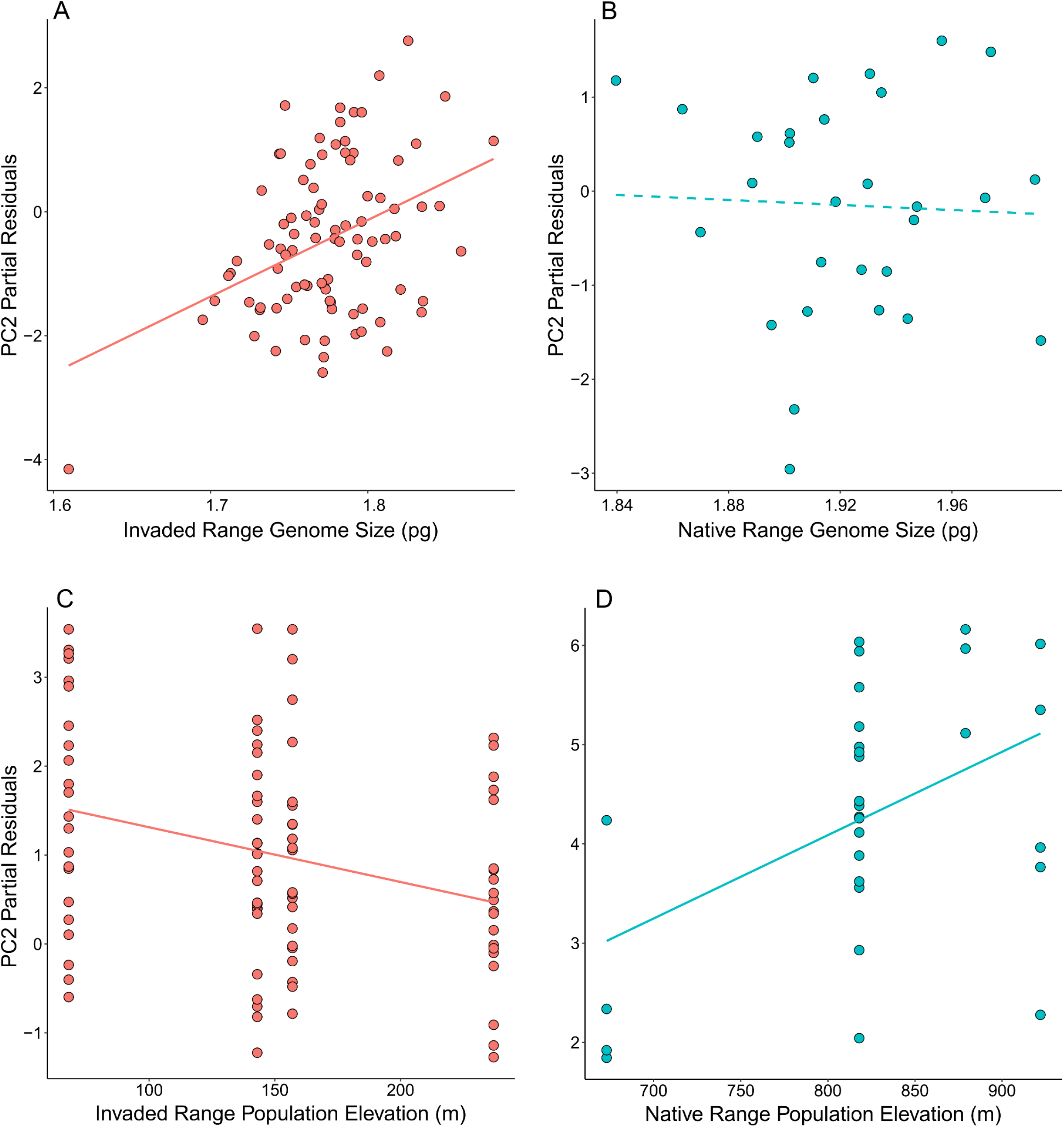
Partial residuals of corrected genome size explaining PC2 variation, plotted separately for the (a) invaded and (b) native ranges. Partial residuals of elevation were also plotted separately for the (c) invaded and (d) native ranges. Solid lines indicate a significant or marginally significant relationship (*p*<0.1), and dashed lines indicate the relationship is not significant.

## Discussion

Although genome size reduction has been proposed as a potential consequence of introduction, the effects of genome size on invasion dynamics and possible contributions to invasibility are not well understood. We surveyed genome size in populations from coastal California, reflecting the oldest areas of introduction in North America, and its native range source populations in Spain, to test for differences across ranges, and its effects on growth and reproduction. We found remarkable variation between and within ranges, with corrected 2C DNA content varying 1.24-fold across invaded and native ranges in California and western Europe, respectively. Larger genomes were associated with later flowering, delayed development and lower elevations only in the invaded range, consistent with selection for faster reproduction at higher elevations favouring smaller genomes. Additionally, the absence of clear relationships in the native range, compared to introduced populations in California, is consistent with adaptation towards reduced genome sizes as a potential contributing factor to its early stages of invasion.

The degree of spread across this set of populations is partially attributable to differences in the distributions of corrected genome size by source range, with a significantly larger mean size from the native range. Our corrected native range estimates are somewhat larger than reported in previously published surveys of YST (Greilhuber 2006; Miskella 2014; Carev *et al*. 2017; Irimia *et al*. 2017). Significant differences in genome size between native and invasive plant populations have been reported in a handful of other systems, such as *Alternanthera philoxeroides* (Chen *et al*. 2011), *Avena barbata* (Crosby et al. 2014), and *Phalaris arundinacea* (Lavergne et al. 2009). To our knowledge, this represents the first documentation of significant differences in genome size by range in this system, as well as the first evidence of support for an advantage to smaller genomes in YST. This observation provides additional evidence to a growing number of systems that suggest selective advantages to smaller genomes in habitats that favour faster growth and development (Achigan-Dako *et al*. 2008; Benor *et al*. 2011; Diez *et al*. 2013; Carta and Peruzzi 2015; Bilinski *et al*. 2018). In addition to genome size differences by range, we also observed substantial variation within individual populations. Intraspecific genome size variation might be of particular importance to invaders, as recently founded populations are likely to be genetically depauperate relative to their source region and, without sufficient variation to adapt to environmental mismatches, may fail to establish and persist (Lee 2002). When standing variation is limited, genome size variation is a possible source of cryptic variation for selection to act upon. Novel stressors encountered by invading populations in their introduced ranges can alter transposable element (TE) dynamics (Grandbastien *et al*. 2005), and Stapley *et al*. (2015) conjecture that resultant TE content variation provides substrate for future adaptation to new environments, especially in the early stages of invasion. Critically, variable rates of TE insertion and deletion are primary contributors to genome size variation (Tenaillon *et al*. 2010), which can exist even in the absence of sequence variation (Jeffrey *et al*. 2016). The extent and frequency of genome size variation that can be generated as a consequence of introduction remains an open question.

Genome size was not a significant predictor of PC1, and we report no evidence for constrained growth due to large genomes in these populations. We found significantly higher mean PC1 scores among invasive genotypes compared to native genotypes, much like previous work (Dlugosch et al 2015). We also found some evidence to support hypothesised constraints of genome size on development-related traits (Benor *et al*. 2011; Carta and Peruzzi 2016; Knight 2005), with larger genomes exhibiting longer time to initiating reproduction and fewer flowers overall (higher PC2 values); however this effect was restricted to invasive populations. Additionally, our linear models indicated faster development associated with the native range (Table 5), despite the observation that native range populations developed slower and produced fewer capitula overall. Our analyses imply an influence of genome size on functional traits that differs by range, in which selection favours smaller genome sizes due to faster development in the invaded range only. The absence of any significant relationship between genome size and development time in the native range suggests genome size emerged as a potential trait under selection only after introduction to the Americas. Novel environmental pressures or demographic changes in the invaded range may have altered how selection acts on genome size variation. However, we cannot say whether possible selection for small genomes was a cause or consequence of invasion, or whether genome size variation conferred a benefit to early introduced YST populations.

We identified a significant interaction between elevation and source range on genome size, in which elevation was negatively correlated with genome size in the invaded range, consistent with predictions of selection for smaller genome sizes where growing seasons are shorter. Again, this correlation was identified only in the invasion, despite no evidence of clinal variation from its source region, where there was no significant association with genome size. In the invaded range, elevation was also significantly negatively correlated with the trait that genome size appears to influence (ie. development time), but had opposing effects on trait variation in the native range. Novel environmental pressures in the invasion associated with early bolting and flowering may underlie this relationship. Additionally, native range populations span elevations (673-922 m) much higher than any of our Bay Area populations (68-237m), suggesting that the benefits of smaller genomes at higher elevations appear context-dependent. Despite some clinal variation, effective population size did not explain genome size in either range. If populations evolved smaller genomes to adapt to shorter growing seasons, we expected selection would be more efficacious when less influenced by the demographics of small populations, and therefore more better at removing larger genome variants. Here, we did not find evidence that founder effects inhibited selection, as we did not observe smaller genomes associated with larger effective populations. This was true within and across the native and invaded ranges, consistent with previous work in this system from Braasch *et al*. (2019) that did not find significant shifts in Ne associated with introduction. It may be worth noting that Ne estimates were calculated by linkage disequilibrium Ne, which relies on sequence variation. Studies of snapping shrimp (Jeffrey *et al*. 2016), prokaryotic taxa (Andreani *et al*. 2017) and alligator weed (Chen *et al*. 2014) suggest that genome size does not predict synonymous sequence variation. If genome size changes offer compensatory variation as targets of selection, it is not expected that Ne estimates could reflect those differences.

Unlike a previous study of YST genome size variation (Irimia *et al*. 2017), we identified significantly larger corrected genome sizes in native range populations, compared to invaded range populations. Critically, we detected an influence of measurement date on genome size estimates, in which later estimates tended to yield smaller genome sizes, and this effect was more pronounced among native range genotypes. Our genome size measurements were taken over five months, during which the plants progressed from rosette formation in early development, through early senescence. Underestimation of nucleus size during flow cytometry can occur when interference from secondary compounds inhibits proper staining of DNA (Doležel *et al*. 2007). Chemical compounds in the cytosol known to inhibit DNA staining vary in concentration, resulting in apparent genome size variation due to stoichiometric errors (Doležel and Bartoš 2005). The magnitude of these errors varies across species (Doležel *et al*. 2007), tissue type (Jedrzejczyk and Sliwinska 2010), developmental stage (Witzell *et al*. 2003; Wam *et al*. 2017), chemical composition in cytosol and standardisation method (Noirot *et al*. 2000; Noirot *et al*. 2003; Noirot *et al*. 2005). Methodological considerations, such as nuclei isolation buffer choice (Loureiro *et al*. 2007) and standardisation protocol (Price *et al*. 2000; Noirot *et al*. 2003), can mitigate some of these errors, but cannot eliminate them entirely, and variation in chemical inhibition of DNA staining persists. Although the identity and function of these compounds are not well-understood in YST, many plant secondary metabolites are produced as a chemical defence against herbivory or pathogen attack (Moore *et al*. 2014). In YST, native range plants experience more complex microbial environments than their invasive counterparts (Lu-Irving *et al*. 2019), and have stronger immune responses during root exposure to bacterial pathogens (Carpenter 2019). We stained nuclei from leaf tissue for flow cytometry, which also varies in chemical composition and allelopathy by range (Irimia *et al*. 2019), with weaker effects in California populations, specifically. Whether heightened chemical defences manifested as less efficacious staining in YST is speculative, and we do not know to what extent staining inhibition explains the interaction effect between range and estimation date. Noirot *et al*. (2003) found a significant effect of estimation date on coffee plant genome sizes, but to the best of our knowledge, very few studies have explicitly addressed or reported it as a significant effect. In YST, we are not aware of any other genome size survey that included discussion of potential effects of estimation date or plant developmental stage on flow cytometry estimates in YST, and the importance of correcting for effects of date across experiments is unclear.

## Acknowledgements

We thank Abreeza Zegeer for assisting with greenhouse experimental set up and management. We thank Paula Campbell and John Finch from the University of Arizona Cytometry Core Facility, and Anthony Baniaga for their assistance preparing flow cytometry protocols for genome size estimation. We would also like to thank Joseph Braasch for supplying effective population size estimates and helpful feedback. We are most grateful to Rhiannon Bauer, Alexus Cazares, Shelby Dolgaard and Alex Skomro for their indispensable help with collecting common garden data. Funding was provided by USDA grant #2015-67013-23000 and NSF grant #1750280 to KMD.

## Author Contributions

FAC and KMD designed the research. FAC collected common garden and flow cytometry data. FAC and KMD conducted data analysis and interpretation. FAC and KMD wrote the manuscript.

