## Supplementary Information for "Ecological effects of genome size in yellow starthistle (*Centaurea solstitialis*) vary between invaded and native ranges"

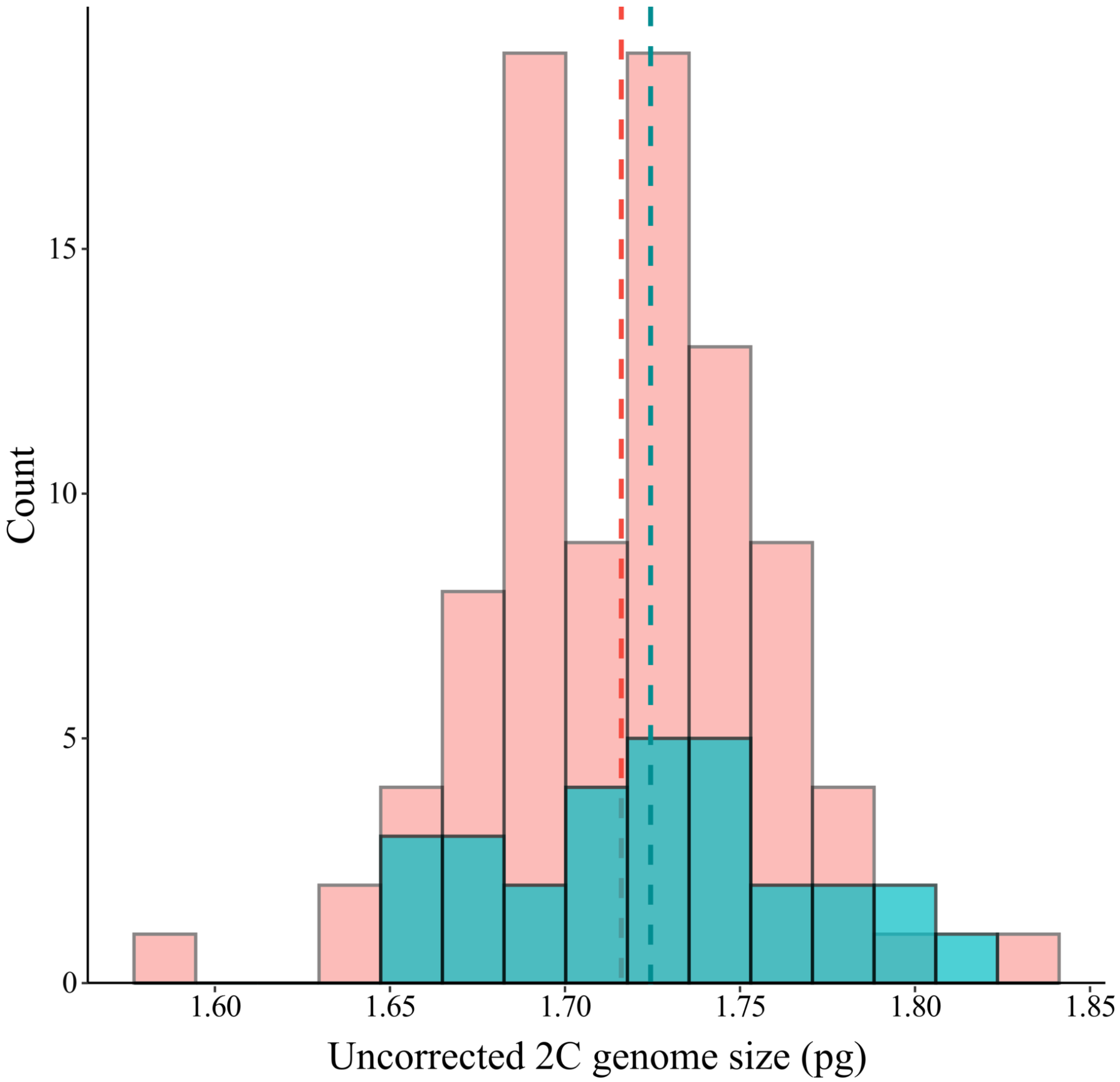


**Figure S1.** Distribution of uncorrected genome sizes by range (blue=native, pink=invaded, N=119) .  Dashed lines are the mean genome sizes within the native (1.716pg) and invaded ranges (1.724pg).
